# Targeting Mycobacterial Transpeptidases: Evaluating the Roles of Ldt and PBP Inhibition in Suppressing *Mycobacterium smegmatis*

**DOI:** 10.1101/2025.01.28.635326

**Authors:** Mariska de Munnik, Karina Calvopiña, Patrick Rabe, Christopher J. Schofield

## Abstract

β-Lactams demonstrate promising in vitro activity against *Mycobacterium* species and are being explored for tuberculosis treatment; however, evidence of their in vivo efficacy versus *Mycobacterium tuberculosis* remains limited. To achieve broad clinically relevant potency, optimisation of the classical β-lactam scaffolds or development of new or non-β-lactam inhibitors for mycobacterial transpeptidases is likely required. In mycobacteria, potential targets of β-lactams include L,D-transpeptidases (Ldts) and penicillin-binding proteins (PBPs). Reports suggest that dual inhibition of Ldts and PBPs may be necessary to achieve effective anti-mycobacterial activity, yet the specific contributions of Ldt and PBP inhibition to the β-lactam antibacterial mechanisms are not understood. We used fluorogenic substrate mimics to investigate the effects of β-lactams and reported Ldt_Mt2_ inhibitors on *Mycobacterium smegmatis* (*Msm*), assessing their impacts on the transpeptidase activities of Ldts and PBPs in living cells. The results reveal a statistically significant correlation between both Ldt and PBP inhibition and *Msm* growth suppression; though under the tested conditions a stronger correlation between Ldt inhibition and *Msm* growth suppression was observed. Notably, inhibition of both PBPs and Ldts was observed in all active inhibitors, though β-lactams manifest increased potency of PBP inhibition. The combination of the β-lactams meropenem and faropenem with selected Ldt_Mt2_ inhibitors manifested an additive inhibitory effect against *Msm*. Our results highlight the importance of further optimising PBPs and Ldt transpeptidase inhibition, particularly of Ldts, to improve β-lactam efficacy versus mycobacteria.

## Introduction

The discovery of β-lactam antibiotics represented a revolution in treatment of infections, and they remain the most important class of antibiotics in clinical use.(1) β-Lactams act as mechanism based inhibitors of the penicillin-binding proteins (PBPs), an essential class of enzymes involved in the biosynthesis of the peptidoglycan layer which is present in every bacterial species, which have D,D-transpeptidase or D,D-carboxypeptidase activities (Figure 1A).(2, 3) The effectivity of β-lactams, including penams, cephalosporins, carbapenems, penems, and monobactams (Figure 1B), is increasingly compromised by resistance, in particular by β-lactamases which are enzymes that catalyse β-lactam hydrolysis (Figure 1A).(1) The presence of a chromosomally encoded Ambler class A β-lactamase (BlaC) has been commonly thought to be a reason why *Mycobacterium tuberculosis* (*Mtb*), the causative agent of tuberculosis (TB) that is estimated to cause 1.6 million deaths per year,(4) is resistant to β-lactams.(5) However, the promising *in vitro* activities of carbapenems in combination with the β-lactamase inhibitor clavulanic acid against *Mtb* have generated interest in reappraisal of β-lactams for TB treatment,(6–9) including with respect to optimisation of cephalosporins for TB treatment. (10–12)

**Figure 1.**
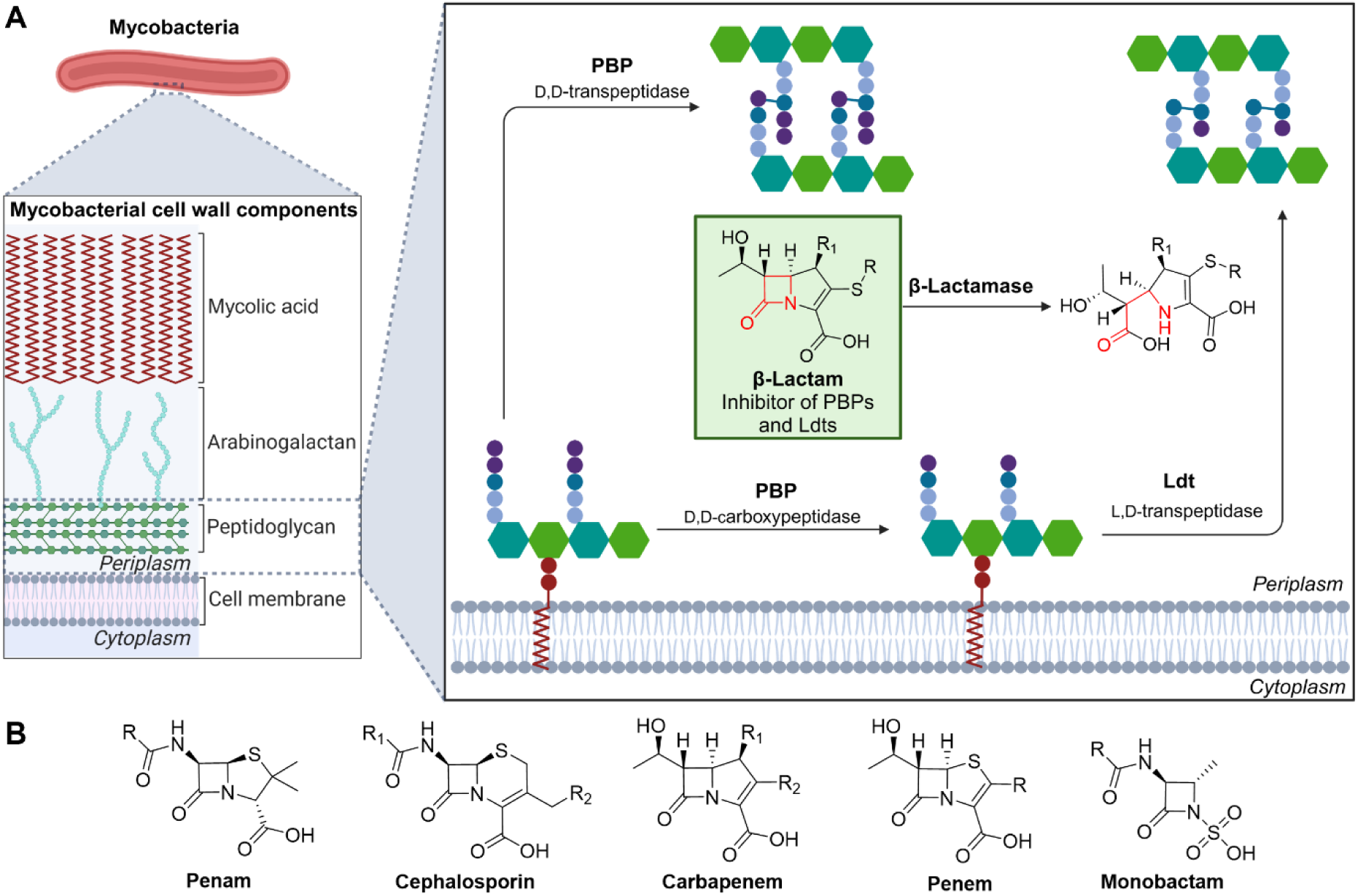
Inhibition of mycobacterial cell wall biosynthesis by β-lactams. **A.** The mycobacterial cell wall is made up of three core components: the mycolic acid, arabinogalactan, and peptidoglycan layers. The final steps of the peptidoglycan biosynthesis include the formation of 3→3 cross-links (catalysed by Ldts) and 4→3 cross-links (catalysed by PBPs), and cleavage of the terminal D-Ala of the pentapeptide monomer to form tetrapeptide monomers (catalysed by PBPs with D,D-carboxypeptidase activity). Cross-links in the peptidoglycan layer of the mycobacterial cell wall can be inhibited by β-lactams (represented here with the core motif of carbapenems); the serine β-lactamases (BlaC in *Mtb*, and BlaS and BlaE in *Msm*) confer resistance. Hexagons in green are N-acetylglucosamine (GlcNAc), hexagons in teal are N-acetylmuramic acid (MurNAc), purple circles are D-Ala, dark blue circles are *m*- DAP, light blue circles are amino acids, dark blue lines are cross-linked amino acids. The figure was created using BioRender.com. **B.** Core motifs of clinically relevant classes of β-lactams.

The activity of β-lactams against *Mtb* is further complicated by the presence of L,D-transpeptidases (Ldts), an important class of enzymes involved in the bacterial cell wall biosynthesis. Ldts are nucleophilic cysteine enzymes that catalyse 3→3 cross-links in the peptidoglycan layer, which are present in exceptionally high quantities in mycobacteria during the stationary phase, compared to the 4→3 cross-links catalysed by the nucleophilic serine PBPs (Figure 1).(13, 14) Several mycobacterial Ldts have been identified as being essential for cell morphology, virulence, aging, and resistance to β-lactams;(15, 16) Ldts are not generally thought to be susceptible to inhibition by most β-lactams, though carbapenems and selected cephalosporins have been observed to inhibit Ldts.(17–20)

*Mtb* contains at least seven PBPs, and five Ldts.(21) It has been suggested that inhibition of Ldt_Mt2_ alone should be sufficient to treat *Mtb*,(22) and several PBPs been identified as being individually essential for growth (PBP3) and/or infection (*e.g.*, PonA1, PonA2, and PBPA).(23, 24) However, there is evidence that inhibition of both Ldts and PBPs may be required for therapeutically relevant inhibition of *Mtb*.(7–9) Dual treatment of amoxicillin/clavulanic acid and meropenem improved inhibition in both *Mtb* and *Mycobacterium smegmatis* (*Msm*).(16, 25) An anecdotal report of a patient treated with the meropenem and amoxicillin/clavulanic acid combination describes the early clearance of *Mtb* from the sputum.(26) However, clinical trials have yielded inconclusive results.(26–28) The available results suggest that for clinically relevant inhibition of *Mtb* via inhibition of PBPs and Ldts, inhibitors tailored towards the *Mtb* transpeptidase targets will be required.

Given the intricate network of cysteine and serine transpeptidases, along with presence of a serine β-lactamase (SBL), the exact modes of action of β-lactams and β-lactam-based inhibitors during mycobacterial chemotherapy are unclear. Recent studies using fluorescent derivatives of β-lactams have been used to address this challenge by identifying and studying the targets of β-lactams in cell lysates of *Mtb*, revealing their binding to various PBPs, Ldts, and BlaC.(29) Pidgeon *et al.* (2019) have also reported on the use of fluorescently labelled substrate mimics of Ldts (TetraRh) and PBPs (PentaFI) to assess Ldt and PBP activity, as well as inhibition by the β-lactams ampicillin and meropenem in whole cells of *Mtb* and *Msm*.(30)

We are interested in further investigating the inhibition of mycobacteria by different classes of β-lactams, as well as selected new types of Ldt_Mt2_ inhibitors (including cephalosporins) recently reported by us.(31, 32) We envisaged that differentiating between Ldt and PBP inhibition, and correlating the extent of transpeptidase inhibition with the inhibition of growth may provide insight that will help enable focussed inhibitor optimisation efforts.

Here we report studies using *Msm*, a well validated model organism for *Mtb*,(33) which contains at least five PBPs and six Ldts, as well as the Ambler class A β-lactamase BlaS,(34–36) to assess the activities of PBP and Ldt_Mt2_ inhibitors, and their ability to inhibit PBPs and Ldts in live *Msm* cells, using PentaFI and TetraRh, respectively. The results reveal a statistically significant correlation between both Ldt and PBP inhibition with suppression of *Msm* growth and imply that a combination of Ldt and PBP inhibition will be beneficial for antibacterial activity, with particular potential for improving Ldt inhibition.

## Results

### Inhibition of *Mycobacterium smegmatis* by β-lactams

We first aimed to assess the activity of different classes of β-lactams against *Msm* (Figure 2, Table S1). Selected carbapenems (meropenem, imipenem, ertapenem, and doripenem; **2**-**5**) manifested potent inhibition of *Msm* (MIC 0.125-4 μg/mL) in the absence of clavulanic acid. This observation is in line with the proposal that carbapenems act as inhibitors of the *Mtb* SBL BlaC.(6, 37, 38) Imipenem (**2**) was particularly potent (MIC 0.125 μg/mL), in accord with reported susceptibility of *Msm* towards imipenem as determined by disc diffusion.(36) The penem faropenem (**1**) also manifested inhibition of *Msm* without the presence of clavulanic acid, though was less potent than the carbapenems (MIC of 16 µg/mL).

**Figure 2.**
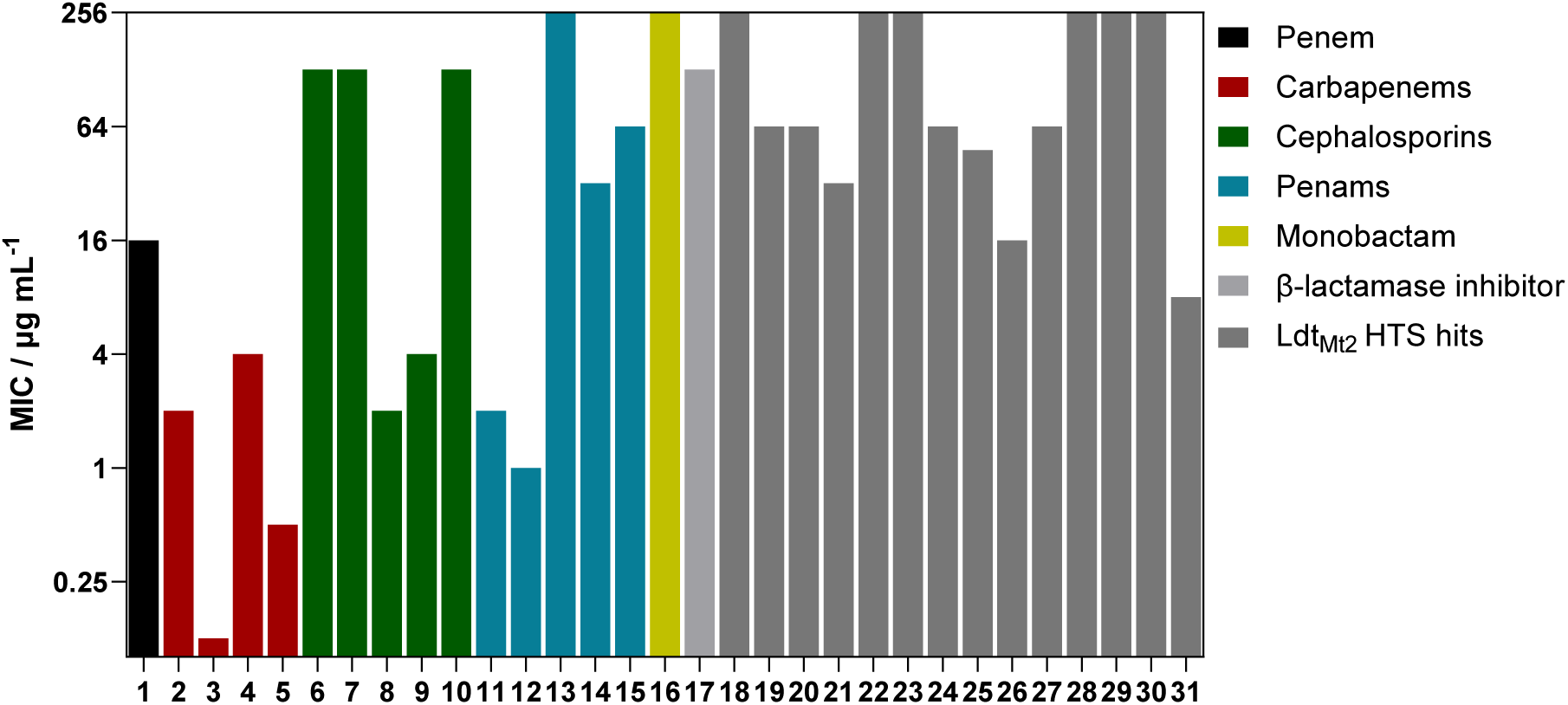
Minimum inhibitory concentrations of 1-31 against *Mycobacterium smegmatis*. The MICs of **1**-**31** against *Msm* were determined with three independent repeats. Note that the highest tested concentration was 128 µg/mL, and inactive compounds are represented here as exceeding an MIC of 128 µg/mL. MIC values and compound structures are given in Table S1.

None of the tested cephalosporins, penams, or the monobactam aztreonam inhibited *Msm* in the absence of clavulanic acid (**6**-**16**; MIC >128 μg/mL), in line with previously reported susceptibility studies as determined by disc diffusion of *Msm*.(36) This observation is likely in part due to β-lactamase activity, because the presence of clavulanic acid (100 µg/mL),(39) which by itself did not potently inhibit *Msm* (MIC 128 µg/mL), significantly improved the activities of ampicillin (**11**) and amoxicillin (**12**) (MICs of 2 µg/mL and 1 µg/mL, respectively).

The activities of ampicillin and amoxicillin versus a ΔblaS strain has been reported (MIC of 16 µg/mL and 1 µg/mL, respectively).(40) In contrast, the inhibitory potencies of oxacillin (**13**), penicillin G (**14**), and carbenicillin (**15**) varied from poor to moderate (MIC >128 µg/mL, 32 µg/mL, and 64 µg/mL, respectively). The presence of clavulanic acid had no impact on the cephalosporins ceftazidime (**6**), ceftriaxone (**7**), and cefepime (**10**) (MIC 128 µg/mL), in contrast to the reported MIC of ceftriazone of 32 μg/mL in the presence of clavulanic acid.(36) However, cephalothin (**8**, MIC 2 µg/mL) and cefmetazole (**9**, MIC 4 µg/mL) showed potent inhibition. Addition of clavulanic acid (100 µg/mL) did not enable inhibition of *Msm* by the monobactam aztreonam (**16**, MIC >128 µg/mL).

To evaluate the potential of various classes of β-lactamase inhibitors in restoring cephalosporin activity, we compared the MICs of cephalothin in the presence of clavulanic acid, tazobactam,(41) sulbactam,(42) BLI-489,(43) and the non β-lactam SBL inhibitors xeruborbactam, a bicyclic boronate and the diazabicyclooctane (DBO) avibactam (Table S2). (44, 45) The addition of sulbactam, xeruborbactam, or avibactam did not enhance the potency of cephalothin. In contrast, tazobactam or BLI-489 (100 µg/mL) increased cephalothin activity to 32 µg/mL and 8 µg/mL, respectively. Clavulanic acid (100 µg/mL) reached the most potent recovery of cephalothin activity (MIC 2 µg/mL). None of the tested β-lactamase inhibitors exhibited inhibitory potency against *Msm* on their own, with MIC values exceeding 128 µg/mL.

### Inhibition of M*ycobacterium smegmatis* by inhibitors of Ldt_Mt2_

A high-throughput screen for *Mtb* Ldt_Mt2_ inhibitors has identified diverse potent electrophilic compounds reacting with the active site nucleophilic cysteine.(31, 32) Despite not being optimised for pharmacokinetic properties, a selection of the inhibitors were active against *Mtb* in macrophages.(31) We set out to investigate the activity of these inhibitors against *Msm*, and additionally tested representative Ldt_Mt2_ inhibitors that did not show activity against *Mtb*, the results of which are summarised in Figure 2 and Table S1. We included the established anti-TB therapeutics ethambutol, isoniazid, and rifampicin as control compounds, which manifested inhibition similar to the reported values (Table S3);(33, 46) note that MIC values reported in the literature for isoniazid vary significantly, and that the Clinical & Laboratory Standards Institute (CLSI) or the European Committee on Antimicrobial Susceptibility Testing (EUCAST) do not provide breakpoints for *Msm*, as it is not a primary clinical pathogen.(47, 48)

The most potent activity against of *Msm* was observed with the sulfonyl pyridine **31**, with an MIC of 8 µg/mL. Interestingly, **31** was also among the most potent compounds tested against *Mtb* (MIC_50_ 11 ± 1.2 μM), though was not active against *Mtb* residing inside the macrophage (MIC_50_ >50 μM). Cyanamide **26** exhibited promising activity with an MIC of 16 µg/mL. Ebsulfur **21** and cyanamide **25** manifested MICs of 32 µg/mL and 48 µg/mL, respectively. Poor activity against *Msm* was observed with **19**, **20**, **24** and **27** (MIC of 64 µg/mL). No inhibition of *Msm* was detected with **18**, **22** and **23** (MIC >128 µg/mL), despite exhibiting moderate inhibition of *Mtb*, while **29**-**30** were inactive versus both *Msm* and *Mtb*.(31)

### Dual treatment of *Mycobacterium smegmatis* with Ldt_Mt2_ inhibitors and β-lactams

To evaluate potential synergy between Ldt_Mt2_ inhibitors and β-lactams, we assessed the effect of dual treatment with the two transpeptidase inhibitor classes. For these studies we selected Ldt_Mt2_ inhibitors with varying efficacy against *Msm* (**19**, **21**, **24**-**26**, and **31**), alongside the inactive compounds **29** and **30** (MIC >128 µg/mL). Meropenem and faropenem were selected as representative β-lactams, based on their inhibitory potency against *Msm* without the presence of clavulanic acid.

We initially assessed effect of Ldt_Mt2_ inhibitors on the MIC of the selected β-lactams in three concentrations (64 µg/mL, 16 µg/mL, and 4 µg/mL; Table S4). The presence of **21**, **25** and **31**, increased the observed activities of both faropenem and meropenem. The increase in activity was apparently more pronounced with the combinations with faropenem. Specifically, the MIC of faropenem decreased from 16 μg/mL to 8 μg/mL, 4 μg/mL, and <0.25 in the presence of 4 μg/mL of **21**, **25**, and **31**, respectively. The MIC of meropenem was only altered from 2 µg/mL to <0.25 µg/mL and 1 µg/mL in the presence of 16 μg/mL **21** and **25**, respectively at a concentration of 16 µg/mL. Further, in the presence of **26** (4 µg/mL), the MIC of faropenem decreased to 8 μg/mL, though **26** (4 µg/mL) did not influence the MIC of meropenem. The presence of **19**, **24**, **29** and **30** had no effect on the MIC of faropenem or meropenem.

Given that compounds **21**, **25** and **31** exhibited increased antibacterial activity in combination with the selected β-lactams, we performed checkerboard analyses of these compounds in combination with faropenem or meropenem (Table 1, Figure S1). We observed increased activity of the Ldt_Mt2_ inhibitor-β-lactam combinations against *Msm* in all cases. According to the recommended interpretation of the determined Fractional Inhibitory Concentration (FIC) indexes of the assessed inhibitor combinations,(49) the results suggest that the combinations of Ldt_Mt2_ inhibitors with faropenem manifest a potentially synergistic effect, with FIC indexes ranging from 0.4 to 0.5, while the combinations with meropenem yielded to an additive effect, with FIC indexes between 0.6 and 0.7 (Table 1, Figure S1).

**Table 1.**
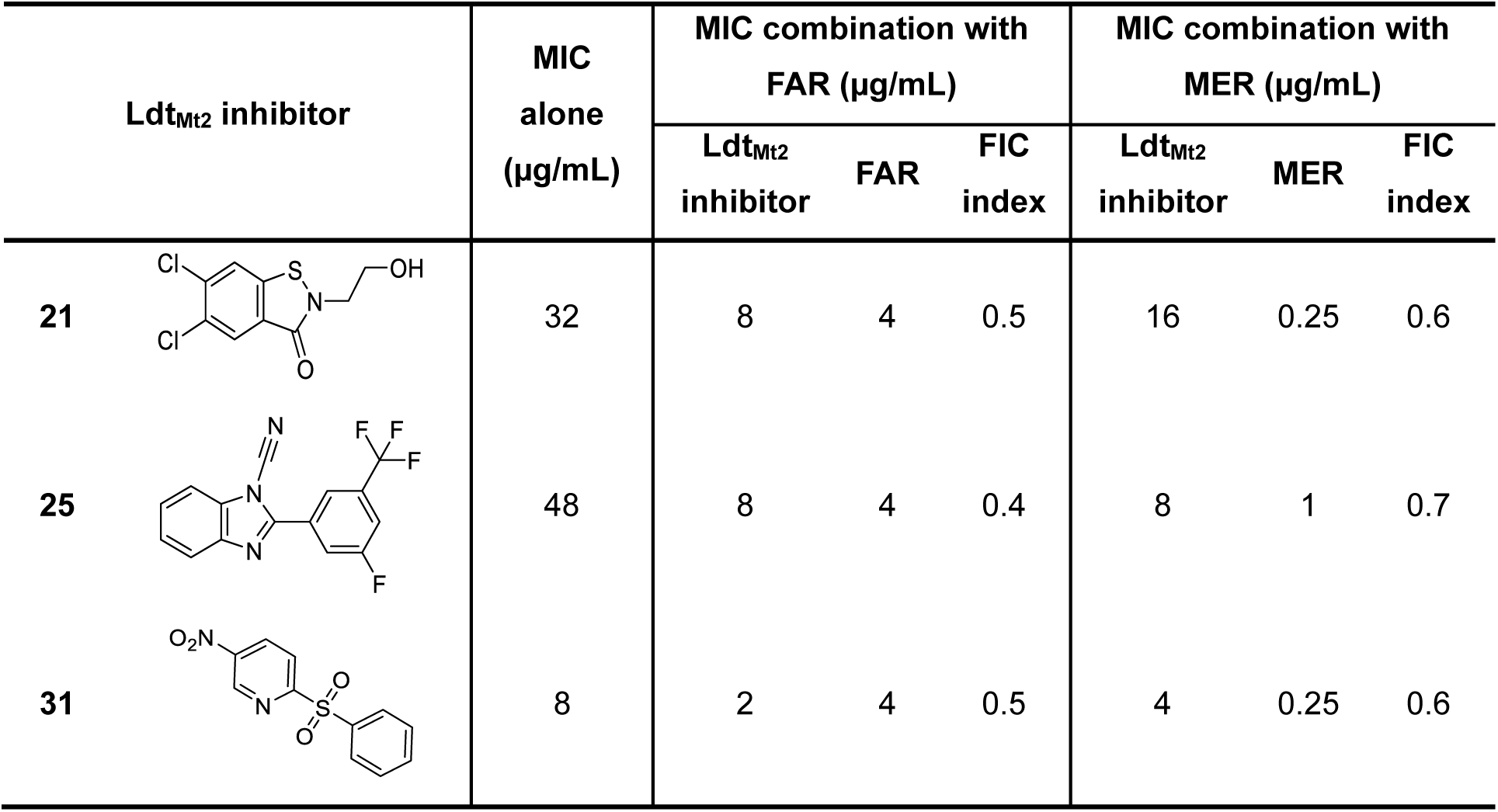
The effects of Ldt_Mt2_ inhibitors 21, 25, and 31 in combination with faropenem or meropenem. FIC index represents the Fractional Inhibitory Concentration index of the indicated combinations of inhibitors, calculated using 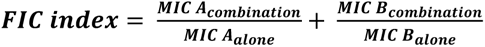.(76) FAR = faropenem (MIC alone of 16 μg/mL), MER = meropenem (MIC alone of 2 μg/mL). The results of all tested combinations are given in Figure S1.

| Ldt<sub>Mt2</sub> inhibitor | MIC alone (µg/mL) | MIC combination with FAR (µg/mL) |  |  | MIC combination with MER (µg/mL) |  |  |
| --- | --- | --- | --- | --- | --- | --- | --- |
|  |  | Ldt<sub>Mt2</sub> inhibitor | FAR | FIC index | Ldt<sub>Mt2</sub> inhibitor | MER | FIC index |
| **21** | 32 | 8 | 4 | 0.5 | 16 | 0.25 | 0.6 |
| **25** | 48 | 8 | 4 | 0.4 | 8 | 1 | 0.7 |
| **31** | 8 | 2 | 4 | 0.5 | 4 | 0.25 | 0.6 |

### Inhibition of the incorporation of fluorescent peptides

To investigate a potential correlation between *Msm* growth inhibition and the inhibition of PBPs and Ldts, we employed the fluorescent probes PentaFI and TetraRh, which mimic the substrates for PBPs and Ldts, respectively (Figure S2), as described by Pidgeon *et al.* (2019).(30) Incubating *Msm* with the inhibitor of interest alongside PentaFI or TetraRh should lead to a dose-dependent reduction in fluorescent signal as a result of inhibition of PBPs or Ldts, respectively (Figure S2), as demonstrated with meropenem and ampicillin with *Msm*.(30)

Optimisation of the fluorescence assays was carried out by assessing the time-dependent incorporation of PentaFI and TetraRh into *Msm*. With 50 µM PentaFI, we observed a time-dependent increase in the fluorescence signal over 24 h, while the negative control, treated with meropenem-clavulanic acid (100 µg/mL), showed no such increase (Figure S3A). Incubation with 50 µM TetraRh exhibited high levels of fluorescent incorporation within the first hour, with minimal subsequent increase. The negative control (100 µg/mL meropenem-clavulanic acid) exhibited reduced fluorescence, but manifested some fluorescence (Figure S3B). When the concentration of TetraRh was reduced to 0.5 µM, a time-dependant increase in the fluorescence signal was observed over 24 h, with the negative controls lacking an apparent fluorescent signal (Figure S3D). As reported,(30) simultaneous incubation of both TetraRh (50 µM) and PentaFI (50 µM) showed only detectable incorporation of TetraRh, likely due to the hight emission intensity of TetraRh.

We performed dose-response studies using this assay to determine pIC_50_ values for cellular inhibition of Ldts and PBPs in *Msm*, focussing on representative β-lactams from different classes and Ldt_Mt2_ inhibitors (Figure 3, Table S1). In most cases, the inhibition of PBPs was more pronounced than Ldt inhibition, with the exceptions being the Ldt_Mt2_ inhibitors **25** and **31**, where a modest dominance of Ldt inhibition was observed (Figure 3A). Amongst all the tested compounds, the carbapenem imipenem displayed the most potent apparent inhibition of both the Ldts and PBPs in *Msm*, with pIC_50_ values of 7.5 ± 0.047 and 9.2 ± 0.054, respectively (Figure 3B,C, Figure S4). Other carbapenems, including meropenem, ertapenem and doripenem, and the penem faropenem, also exhibited potent inhibition of transpeptidase activity of PBPs (pIC_50_ values ranging from 5.7 to 6.6) and Ldts (pIC_50_ values ranging from 5.1 to 6.1).

**Figure 3.**
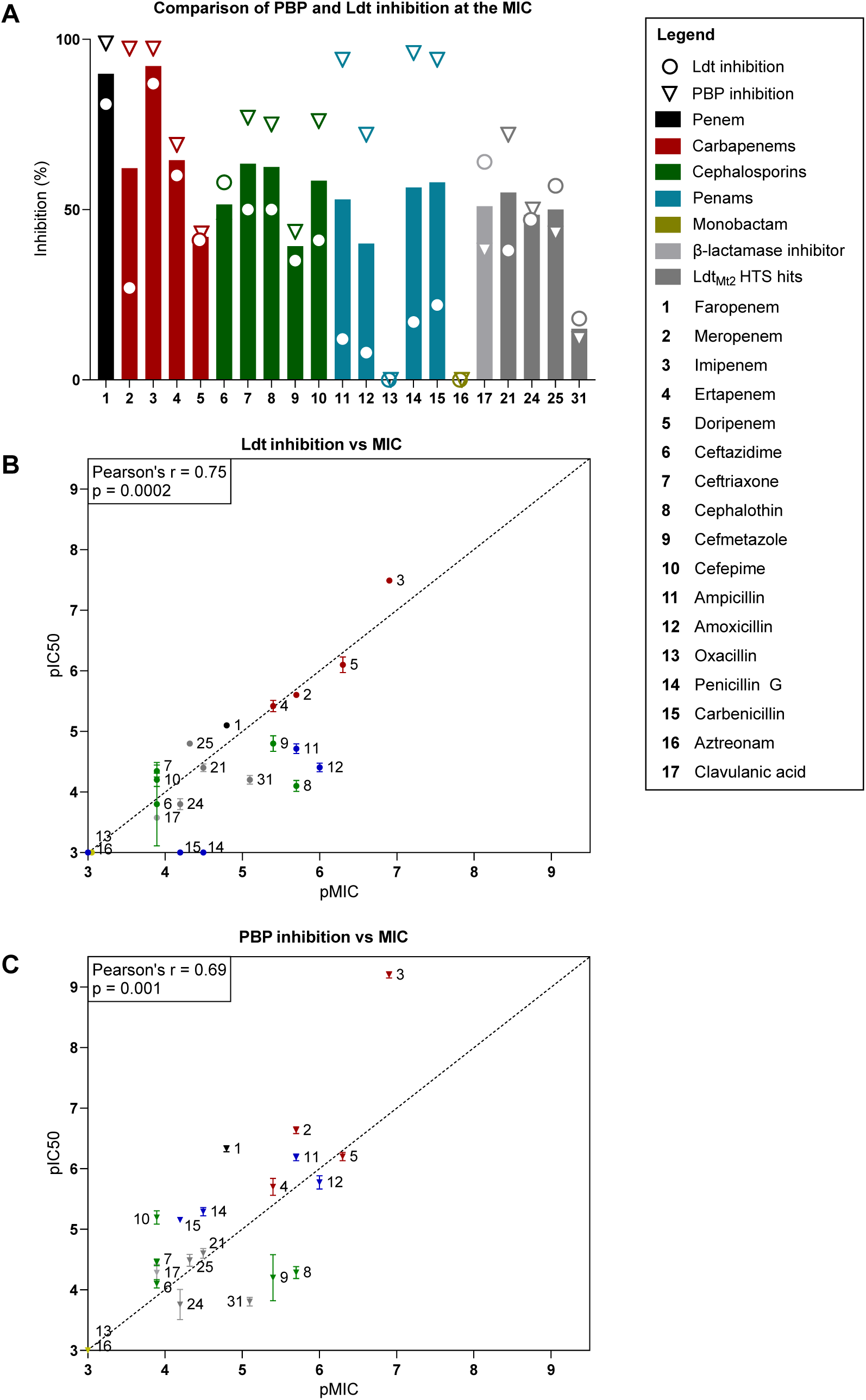
Fluorescent peptide incorporation assay investigating the inhibition of Ldts and PBPs in live *Mycobacterium smegmatis* cells. **A.** The inhibition of the PBPs and the Ldts at the MIC of the respective inhibitors was investigated. Bars represent the average between Ldt and PBP inhibition. **B**. The inhibitory potencies (pIC_50_’s) of the inhibitors for *Msm* Ldts were determined using the dose-response assays with TetraRh, and plotted against the *Msm* pMIC. **C.** The inhibitory potencies (pIC_50_’s) of the inhibitors for *Msm* PBPs were determined using the dose-response assays with PentaFI, and plotted against the *Msm* pMIC. Note that pMIC values arise from logarithmic transformation of MIC (in g/mL) and that pIC_50_ values arise from logarithmic transformation of IC_50_ (in g/mL). Data points represent mean and error bars represent standard deviation. Values and compound structures are given in Table S1.

In contrast, all the assessed cephalosporins (ceftazidime, ceftriaxone, cephalothin, cefmetazole, and cefepime) showed limited inhibition of TetraRh or PentaFI incorporation, despite the presence of clavulanic acid (100 µM). PBP inhibition ranged between pIC_50_ values of 4.1 ± 0.069 for ceftazidime and 5.2 ± 0.11 for cefepime, while Ldt inhibition ranged between pIC_50_ of 3.8 ± 0.69 for ceftazidime and 4.8 ± 0.13 for cefmetazole. The penams ampicillin, amoxicillin, penicillin G, and carbenicillin exhibited moderate to potent inhibition of PentaFI incorporation in the presence of clavulanic acid (100 µM), with the pIC_50_ values ranging between 5.2 ± 0.042 for carbenicillin and 6.2 ± 0.056 for ampicillin. However, inhibition of TetraRh incorporation was very limited, with the most potent Ldt inhibition being a pIC_50_ value of 4.7 ± 0.079 for ampicillin. Oxacillin and the monobactam aztreonam, which showed no inhibition of growth of *Msm*, did not apparently inhibit either Ldts or PBPs.

The inhibitors identified in the HTS against Ldt_Mt2_ (**21**, **25**, and **31**) demonstrated limited inhibition of TetraRh incorporation, with pIC_50_ values ranging between 3.8 and 4.8, correlating with their relatively poor MIC values. Interestingly, these compounds also acted as PBP inhibitors of similar potency (pIC_50_ values ranging between 3.7 and 4.6), despite being identified as Ldt inhibitors. In contrast, **31**, which manifested a significantly lower MIC of 8 µg/mL, showed only limited Ldt inhibition (pIC_50_ value 4.2 ± 0.071), and no PBP inhibition, suggesting a potentially alternative mechanism of action.

When evaluating the inhibition of PentaFI and TetraRh incorporation at inhibitor concentrations corresponding to the MIC, combined PBP and Ldt transpeptidase inhibition varied in most cases from 64% with ertapenem to ∼40% with cefmetazole, amoxicillin, and doripenem (Figure 3A). Outliers to this observation are the ∼90% inhibition of PBP and Ldt transpeptidase activity in the cases of imipenem and faropenem, and 15% inhibition in the case of the sulfonyl pyridine derivative **31**.

Apparently dual inhibition of PBPs and Ldts was observed with all the β-lactam inhibitors, except for the penams penicillin G and carbenicillin. Additionally, ampicillin and amoxicillin manifested considerably lower Ldt inhibition compared to PBP inhibition. We observed a relatively strong correlation between Ldt inhibition and *Msm* MIC (Figure 3B); Pearson correlation analysis revealed a significant positive correlation (r = 0.75, p < 0.001, n = 19), though the correlation was stronger on exclusion of the penams (r = 0.83, p < 0.001, n = 15). A weaker, but significant correlation was observed between PBP inhibition and MIC (r = 0.69, p < 0.05, n = 19; Figure 3C).

### Biochemical inhibition of the *Mycobacterium tuberculosis* PBP3

Following the observation that Ldt_Mt2_ inhibitors, in particular the ebsulfur derivative **21** (pIC_50_ 4.6 ± 0.080), apparently inhibit PBPs in a cellular context, we investigated the inhibition of the essential isolated recombinant PBP3 of *Mtb* by **18**-**31** (Figure 4A, Table S1, Figure S5).(23, 50, 51) Interestingly, despite cellular inhibition of *Msm* PBPs, most of the assessed compounds manifested limited inhibitory potency (pIC_50_ ≤ 4.5 for **19**-**21** and **26**) or no significant inhibition (pIC_50_ < 3.4 for **18**, **22**, **25**, and **28**-**31**) against PBP3. The exception was cephalosporin **24** (pIC_50_ of 6.1 ± 0.089). Notably, **24** has been identified as an inhibitor of both Ldt_Mt2_ and BlaC, although it does not display potent anti-TB activity,(31) or significant anti-*Msm* activity (MIC 64 μg/mL).

**Figure 4.**
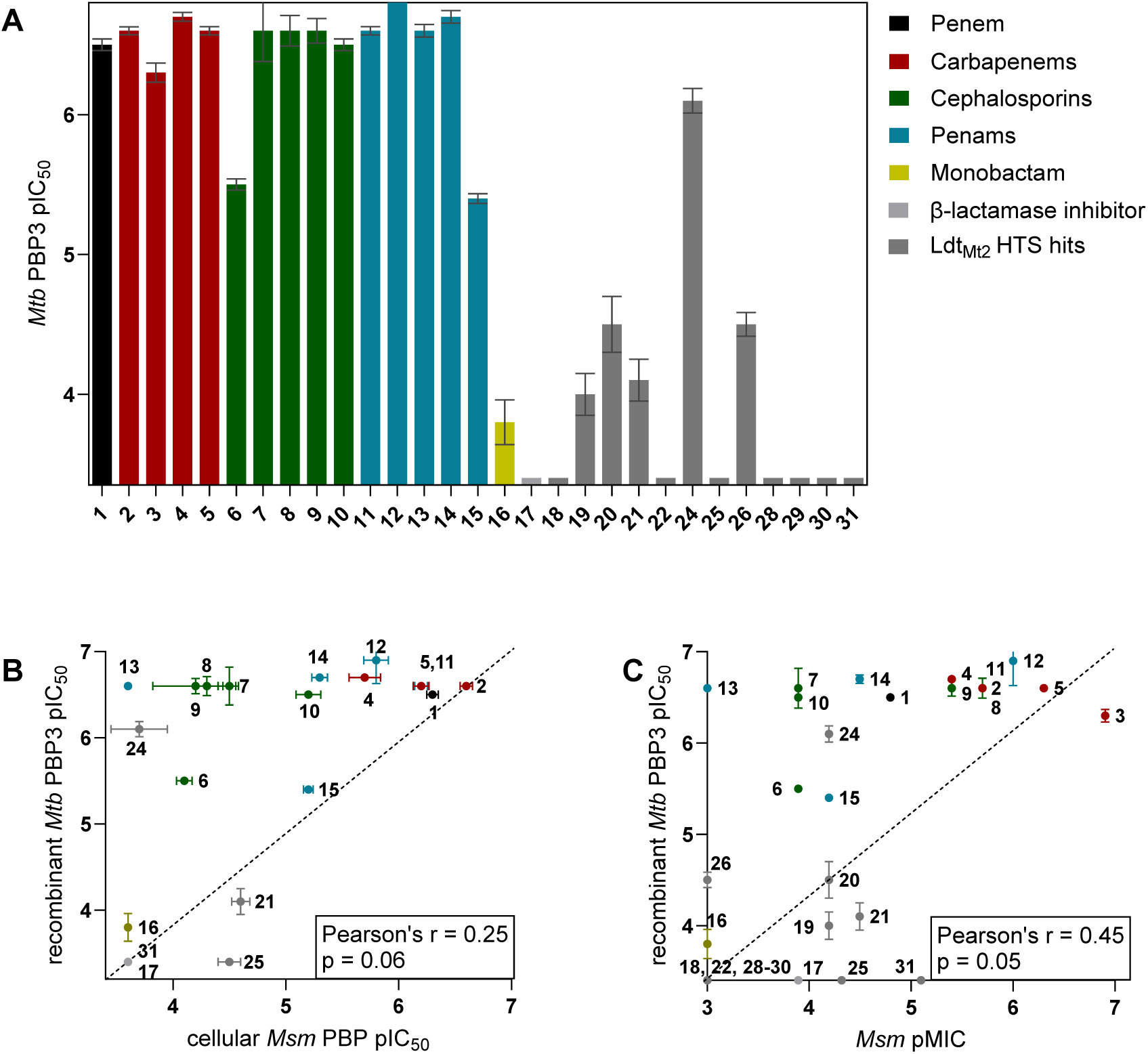
Inhibition of *Mycobacterium tuberculosis* PBP3 by 1-31. **A.** The inhibitory potencies (pIC_50_’s) of the inhibitors for *Msm* Ldts were determined using the dose-response assays using the fluorometric S2d assay,(72–74) applying the procedure optimised for *P. aeruginosa* PBP3.(75) **B.** The *Mtb* PBP3 pIC_50_ values did not show a significant correlation with the *Msm* PBP inhibition determined in the fluorescent peptide incorporation assays. **C.** The *Mtb* PBP3 pIC_50_ values did not show a strong correlation with the MIC’s of *Msm*. Values and compound structures are given in Table S1, and data curves are given in Figure S5.

We also evaluated inhibition of isolated *Mtb* PBP3 by β-lactams (Figure 4A, Table S1, Figure S5). Most β-lactams potently inhibited PBP3, with pIC_50_ values approaching, or reaching, the upper assay limit (pIC_50_ of 6.8). Ceftazidime, known for its potent inhibition of PBP3 in Gram-negative bacteria,(52) and carbenicillin displayed slightly lower potencies (pIC_50_ values of 5.5 and 5.4, respectively). Additionally, aztreonam, which is effective against PBP3 in Gram-negative but not Gram-positive bacteria,(53) manifested very low inhibition of *Mtb* PBP3 with a pIC_50_ value of 3.8.

No significant correlation was observed between the cellular inhibition of the PBPs of *Msm* by **1**-**31** and the inhibition of *Mtb* PBP3 under current assays conditions as evidenced by Pearson’s r analysis (r = 0.25, p > 0.05; Figure 4B). Only weak correlation between the MIC against *Msm* and the inhibition of *Mtb* PBP3 was observed (r = 0.45, p = 0.05; Figure 4C).

### Sulfonyl pyridines exhibit promising anti-mycobacterial activity

Of the inhibitors identified in the HTS against *Mtb* Ldt_Mt2_,(31) the sulfonyl pyridine **31** manifested the most potent inhibition of *Msm*. However, the fluorescent peptide incorporation assays revealed only limited evidence for Ldt and PBP inhibition (Figure 3), suggesting a potential alternative mechanism of action. However, given its notable activity, we evaluated a series of derivatives of **31** against *Msm* (Figure 5, Table S5).

**Figure 5.**
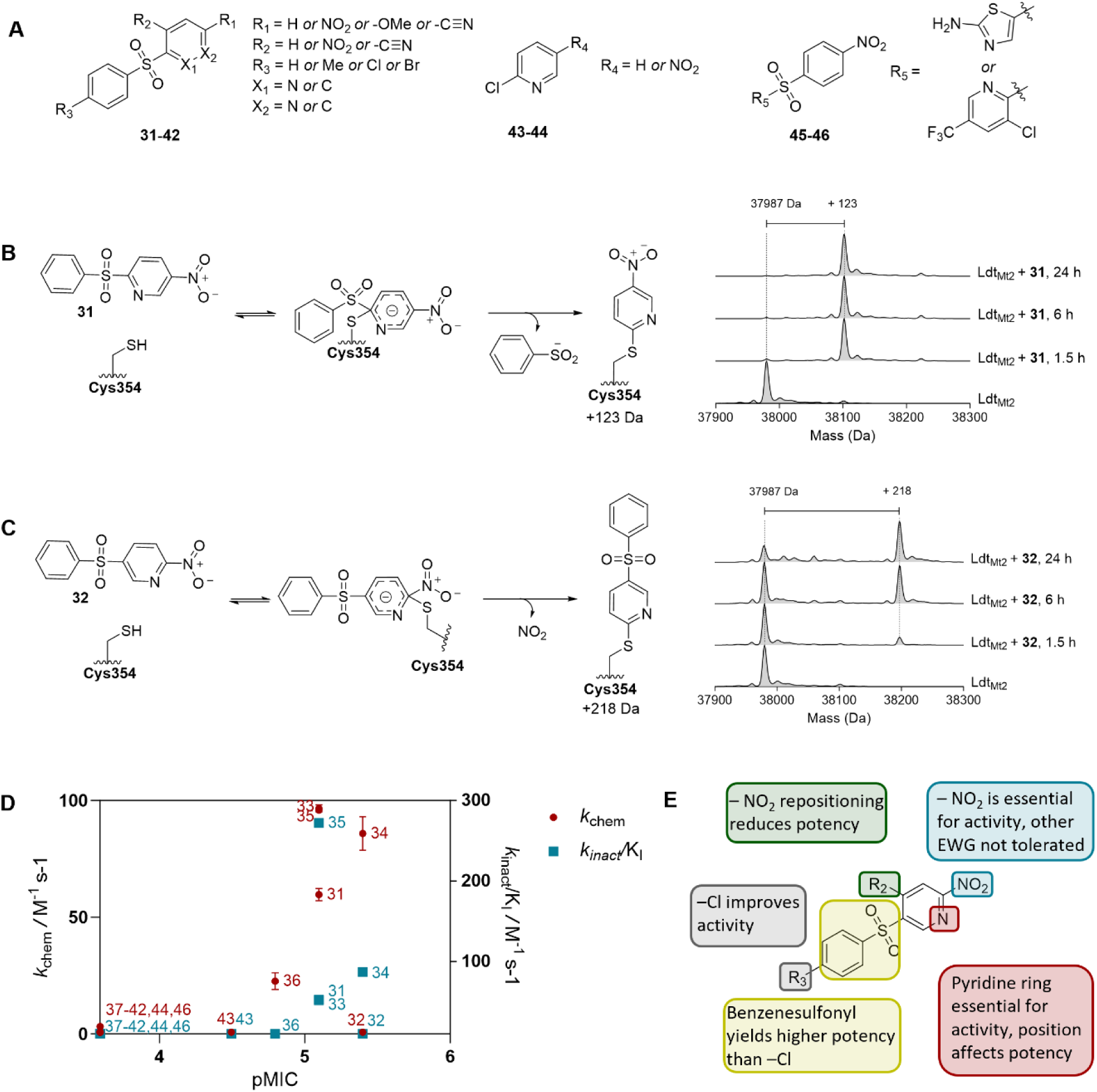
Sulfonyl pyridines are inhibitors of *Mycobacterium smegmatis*. **A.** Sulfonyl pyridines **31**- **46** were tested against *Msm*. Full compound structures and MICs are given in Table S5. **B.** Mass spectrometry studies suggest selected sulfonyl pyridines to react with the nucleophilic cysteine of Ldt_Mt2_ via nucleophilic aromatic substitution, as exemplified with **31**. **C.** Mass spectrometry studies with **32** suggest that repositioning of the nitrogen atom on the pyridine ring reduces reactivity with Ldt_Mt2_ and alters the leaving group. **D.** The MIC against *Msm* does not correlate with the rate of inhibition against Ldt_Mt2_ (*k*_inact_/K_I_, in red) or with the rate of intrinsic thiol reactivity (*k*_chem_, in blue). **E.** Findings of the structure-activity relationship studies with **31-46** based on the MIC against *Msm*.

For derivative **32**, altering the position of the nitrogen atom in the pyridine ring reduced the MIC to 4 µg/mL. Modifications to the phenyl ring, such as addition of a methyl group (**33**) or bromide (**35**), did not change the MIC, while substitution with a chloride (**34**) decreased the MIC to 4 µg/mL. Moving of the nitro group from the *para* to the *ortho* position (**36**) increased the MIC to 16 µg/mL. Notably, the removal of the nitro group entirely abolished activity against *Msm*, regardless of the position of the nitrogen atom on the pyridine ring (**37**-**39**), or substitution of the nitro group with a methoxy group (**40**) or a nitrile in *para* (**41**) or *ortho* (**42**) positions. Substituting the phenylsulfonyl group with a chloride increased the MIC to 32 µg/mL (**43**). Similarly to **37**-**42**, elimination of the nitro group in that moiety resulted in the loss of activity against *Msm* (**44**). Furthermore, nitrophenyl sulfonyl derivatives **45** and **46** did not exhibit activity against *Msm*.

While Ldts may well not be the (only) cellular target of the sulfonyl pyridines, we examined their reactivity with the nucleophilic thiol of Ldt_Mt2_. Protein-observed solid-phase extraction MS (SPE-MS) assays with Ldt_Mt2_ showed that **31**, **33**-**36**, **41**, **42** and **46** all reacted with Ldt_Mt2_ through nucleophilic aromatic substitution of the sulfonyl group on the pyridine ring (Figure 5B, Table S5, Figure S9). In contrast, the mass shift observed with **32** corresponds to nucleophilic substitution at the nitro group rather than the sulfonyl group (Figure 5C). Derivatives **37**-**40**, **43** and **44** did not react with Ldt_Mt2_; of these only **43** exhibited moderate activity against *Msm* (MIC 32 µg/mL). Kinetic studies of the reaction with Ldt_Mt2_ (Table S5, Figure S6, S7), and with L-glutathione (GSH; Table S5, Figure S8) were performed to profile the electrophilic reactivity of this class of inhibitors. We observed no correlation between reactivity with Ldt_Mt2_ or GSH and activity against *Msm*; for example, **32** demonstrated significantly lower reactivity with both Ldt_Mt2_ and GSH compared to **31** and **33**-**35**, yet showed increased activity against *Msm* (Figure 5D).

## Discussion

Historically, β-lactams have been considered to be ineffective for treating TB. However, emerging *in vitro* studies demonstrating promising activity against *Mtb* have led to a re-evaluation of this paradigm.(7–9) Nevertheless, clinical studies have yielded inconclusive results regarding the utility of β-lactam antibiotics for the treatment of TB,(26, 28, 54) suggesting that optimisation of inhibitors specifically tailored for *Mtb* may be required. Due to the high level of 3→3 cross-links in the peptidoglycan layer catalysed by the Ldts, Ldts are considered a promising target for the development of inhibitors. Some evidence, however, indicates that effective inhibition may require targeting both Ldts and PBPs.(16) A deeper understanding of the mechanism of action of β-lactams will facilitate targeted efforts these compounds and promote the development of non-β-lactam inhibitors against *Mtb*.

Our results reveal a correlation between MIC values and the extent of inhibition of both PBP and Ldt transpeptidase activities, though a stronger correlation between Ldt inhibition and MIC was observed. Importantly, we observed dual inhibition of PBPs and Ldts for all the tested inhibitors of *Msm*. However, we noted multiple instances (e.g., with meropenem, imipenem, faropenem, ampicillin, penicillin G, and amoxicillin) where complete inhibition of cellular PBP transpeptidase activity did not correspond to growth inhibition. These findings support the importance of Ldts as an important target for *Mycobacterium* spp. inhibition, particularly in combination with PBP inhibition, at least under the tested conditions. They suggest that optimising β-lactams against *Mycobacterium* spp. should focus on enhancing their potency against Ldts, as well as PBPs. However, to better define the relationship between Ldt inhibition and MIC independent PBP inhibition, the development of Ldt-specific inhibitors is of considerable interest.

Carbapenems and penems are known for their ability to inhibit both Ldts and PBPs.(55–58) Correspondingly, we observed potent inhibition of both classes of transpeptidases, particularly by imipenem in the fluorescence based *Msm* cell assays. While faropenem has been identified as the most potent inhibitor of Ldts in recombinant enzyme assays,(18, 59, 60) our findings indicate lower inhibitory potency of faropenem against the Ldts in live *Msm* cells, in comparison to the tested carbapenems. Interestingly, the penams ampicillin and amoxicillin exhibit significant inhibition of *Msm* in the presence of clavulanic acid, correlating with PBP transpeptidase inhibition, but showing much less inhibition of Ldts.

While we observed that **21** and **25** appear to inhibit both Ldts and PBPs in *Msm*, **31** appears to operate via an alternative mechanism of action. Notably, all three compounds manifested similar levels of synergistic or additive inhibition when combined with faropenem or meropenem. Further research is warranted to elucidate the mechanism behind the benefit of dual treatment of β-lactams and **31**, while in the cases of **21** and **25** the increased inhibition of both Ldts and PBPs in the presence of β-lactams is likely to play a role. However, both **21** and **25** contain a reactive electrophilic group, and may act as relatively non-specific inhibitors.

Though the cellular activity of sulfonyl pyridine **31** could not be correlated to the extent of inhibition of PBPs or Ldts, **31** displayed unexpectedly potent activity against *Msm*, likely involving another or an additional mechanism of action. Inhibitory studies with derivatives of **31** suggest that the nitropyridine sulfonyl moiety is critical for its inhibitory activity against *Msm*, as removal of the nitro group or substitution with other electron withdrawing groups, as well as substitution of the pyridine ring for a phenyl ring, abolished activity. Reactivity with nucleophilic cysteine residues (as evidenced by interactions with Ldt_Mt2_ and GSH) did not correlate with activity against *Msm*. Therefore, identification of the mechanism of action of the sulfonyl pyridine compounds is of interest.

Following on the observation that Ldt_Mt2_ inhibitors were able to inhibit both Ldts and PBPs in *Msm*, we assessed the biochemical inhibition of the essential PBP3 of *Mtb*. Interestingly, the extent of inhibition of PBP3 by the Ldt_Mt2_ inhibitors was limited. In contrast, most tested β-lactams demonstrated potent inhibition of *Mtb* PBP3. An exception was aztreonam, which has been shown to interact with *Mtb* PBP3 in crystallographic studies.(61) Aztreonam is known as a potent broad spectrum inhibitor of Gram-negative bacteria, but is less effective against Gram-positive bacteria.(53) *Mtb*, as an acid-fast bacillus, falls in neither these categories, though they can be considered a subclass of Gram-positive bacteria.(62) Overall, using the current assays we found no clear correlation between cellular inhibition of the PBPs and Ldts of *Msm* and the inhibition of recombinant *Mtb* PBP3 and Ldt_Mt2_, though all cellularly active inhibitors were also inhibitors of isolated *Mtb* PBP3, with the exception of **25**.

Our studies utilised *Msm* as a validated model organism for *Mtb*.(33) LdtC is reported to be the main functional Ldt of *Msm*,(63) as well as the main contributor to TetraRh incorporation.(30) LdtC is a closer homologue of *Mtb* Ldt_Mt5_ than Ldt_Mt2_, the latter of which is the main Ldt of *Mtb*, which is a close homologue of *Msm* LdtB.(60) Superimposition of a Ldt_Mt2_ structure with a structural prediction model of LdtC created with AlphaFold,(64) manifests high structural similarity of the two active site regions (Cα RMSD 0.75 Å) and of key residues (Figure S10A). PBP3 from *Msm* and *Mtb* share ∼79% sequence similarity, and manifest structural similarity of their active site domains (RMSD 0.48 Å; Figure S10B).(61) It has yet to be determined which PBPs in *Msm* contribute to the incorporation of PentaFI into the cell wall. However, considering the nature of the assay, it can be inferred likely that only PBPs with D,D-transpeptidase activity are involved, and not those with D,D-carboxypeptidase activity.

The primary β-lactamase of *Msm*, BlaS, has ∼40% sequence similarity and high structural homology with BlaC, the major β-lactamase of *Mtb*, and exhibits particularly efficient penicillinase and cephalosporinase activity.(36) Similarly to BlaC, BlaS can be inhibited by clavulanic acid.(36) However, *Msm* also expresses an additional cephalosporinase, BlaE, which is less sensitive to clavulanic acid inhibition,(36) knowledge consistent with the resistance to several cephalosporins (e.g. ceftazidime and ceftriaxone) observed in our experiments, despite the presence of clavulanic acid.

## Conclusion

Certain β-lactams exhibit promising activity against *Mycobacterium* spp. though optimisation will likely be required to obtain clinically relevant potency. Relevant targets for β-lactams in *Mycobacterium spp*. include the PBPs and the Ldts, with emerging evidence suggesting that dual inhibition of these targets may be essential for optimised antibacterial efficacy, via transpeptidase inhibition. In living *Msm* cells, β-lactams (particularly the carbapenems, penems, and penams) are potent inhibitors of PBPs with transpeptidase activity, reinforcing their potential as therapeutic options. Notably, our results reveal a strong correlation between Ldt inhibition and MIC values, implying the critical role of Ldt inhibition in the inhibition of *Msm*. In addition, we identified the sulfonyl pyridines as promising inhibitors of *Msm*, though they could not be related to inhibition of transpeptidase activity of Ldts or PBPs. Overall, our results suggest that future efforts to optimise β-lactams, alongside non-β-lactam inhibitors of transpeptidases, that focus on enhancing potency against Ldts, may lead to effective inhibition of *Mycobacterium* spp.

## Methods

### Materials

S2d and 2-(6-(((2,4-dinitrophenyl)sulfonyl)oxy)-3-oxo-3H-xanthen-9-yl)benzoic acid (Probe 1) were synthesised as described.(65–67) **18**-**31** were obtained from the GSK HTS compound library. **32**, **36**, and **38**-**42** were from Cortex Organics, **37** and Fmoc-D-iGln-OH were from AmBeed, **33**-**35** and **46** were from Key Organics, **44** and **45** were purchased from Enamine, and 5(6)-TAMRA was from MedChemExpress. Faropenem was purchased from Fluorochem Ltd, meropenem was from Glentham Life Sciences, ceftazidime was purchased from TOKU-E, ceftriaxone and aztreonam were from Molekula Ltd, ampicillin was from Apollo Scientific, and ampicillin was from Alfa Aesar. Isoniazid, ethambutol, and rifampicin were from Cambridge Bioscience Ltd; all other compounds were purchased from Merck.

### Synthesis of fluorescent peptides

PentaFI (D-Ala-D-Ala-L-Lys-D-iGln-L-Ala-Fluorescein) and TetraRh (D-Ala-L-Lys-D-iGln-L-Ala-Rhodamine) were synthesised by solid phase peptide synthesis using a Liberty Blue Automated Microwave Peptide Synthesiser, following stepwise coupling reactions to D-Alanine-Wang resin (715 mg, 0.5 mmol). The respective amino acids D-Ala (1.5 mmol, 3 eq.), L-Lys (2.5 mmol, 5 eq.), D-iGln (2.5 mmol, 5 eq.), and L-Ala (2.5 mmol, 5 eq.), were coupled using the standard procedure with N,N-diisopropylethylamine (DIPEA) in ethyl cyanohydroxyiminoacetate (Oxyma, 1 M; DIPEA/Oxyma 1.5 % (*v*/*v*)) and 7.8% (*v*/*v*) N,N′-diisopropylcarbodiimide (DIC) in DMF. After each coupling step, the resin was washed with DMF and the Fmoc group was deprotected with 20% (*v*/*v*) piperidine in DMF using the standard deprotection procedure. The fluorescent tags 5(6)-carboxyfluorescein (2 eq) and 5(6)-TAMRA (2 eq) were coupled manually, using hexafluorophosphate benzotriazole tetramethyl uronium (HBTU; 2 eq) and DIPEA (6 eq). The resin bound TetraRh and PentaFI were washed with dichloromethane (DCM), methanol and DCM, and treated with a 5 mL solution of 2:1 TFA/DCM (2 h, rt). Solvents were removed by evaporation *in vacuo* and purified by HPLC on a C_18_ column (5 µm, 10 x 150 mm; SunFire, Waters) at a flowrate of 3 mL/min with a gradient of 98% (*v*/*v*) buffer A (0.1% (*v*/*v*) formic acid in H_2_O) and 2% buffer B (0.1% (*v*/*v*) formic acid in acetonitrile) to 50% buffer A and 50% buffer B over 22 min. TetraRh and PentaFI eluted after 18.8 min and 22.1 min, respectively. TetraRh and PentaFI were characterised by ^1^H as well as two-dimensional NMR experiments including ^1^H,^1^H-COSY, ^1^H,^13^C-HSQC and ^1^H,^13^C-HMBC (Figure S12-13, Table S6-7).

### Recombinant protein production and purification

Ldt_Mt2_ was produced in *Escherichia coli* and purified (>95% purity by SDS-PAGE analysis) as previously described.(68)

A codon-optimised synthetic gene (GeneArt, Thermo Fisher Scientific) encoding for *Mtb* PBP3 Δ1-122 was amplified and cloned into the expression vector pCold using Sal1-HF (New England BioLabs) and Not1-HF (New England BioLabs) digestion and ligation using T4 DNA ligase (New England BioLabs) according to the manufacturer’s protocol. The ampicillin resistance gene of the vector was exchanged for the kanamycin resistance gene using Gibson Assembly.(69)

A culture of *E. coli* BL21(DE3) pCold-PBP3 Δ1-122 was grown at 37 °C at 180 rpm in 2xYT media (with 50 µg/mL kanamycin) to OD_600_ of 0.6. Isopropyl β-D-thiogalactopyranoside (IPTG) (0.5 mM) was then added and the culture was incubated at 15 °C at 180 rpm for an additional 16 h. Cells were collected by centrifugation (11,000 x g, 8 min), and stored at −80 °C. The cell pellet was resuspended in HisTrap Buffer A (50 mM Tris-HCl pH 7.5, 500 mM NaCl, 20 mM imidazole) in the presence of DNase I, and lysed using a Continuous Flow Cell Disruptor (Constant Systems, 25 kpsi). The lysates were centrifuged (32,000 x g, 20 min), passed through a 0.45 µm filter, and loaded onto a 5 mL HisTrap column (GE Life Sciences) that had been pre-equilibrated in HisTrap Buffer A. The column was washed with HisTrap Buffer A, followed by a gradient running from 0 % to 100 % (*v/v*) HisTrap Buffer B (50 mM Tris-HCl pH 7.5, 500 mM NaCl, 500 mM imidazole). Fractions containing PBP3 (as observed by SDS-PAGE) were combined, the buffer was exchanged to HisTrap Buffer A, and the HisTag was cleaved using recombinant 3C protease at 4 °C, over 12 h. The HisTag cleaved PBP3 was passed through a 5 mL HisTrap column (GE Life Sciences) and subsequently loaded onto a 300 mL Superdex 75 column (GE Life Sciences) pre-equilibrated in gel filtration buffer (50 mM Tris-HCl pH 7.5, 500 mM NaCl), and eluted over 1 column volume in gel filtration buffer. The identity and purity of PBP3 was confirmed by mass spectrometry (calculated mass 60148 Da, observed deconvoluted mass 60152 Da) and SDS-PAGE (>95% purity).

### Bacterial strain and growth conditions

All experiments using *Msm* refer to strain ATCC 607. *Msm* was grown on Columbia blood agar supplemented with 5% sheep blood at 37 °C. Liquid cultures of *Msm* were grown in Middlebrook 7H9 broth supplemented with 0.1% (*v*/*v*) Tween-80, 0.5 % (*w*/*v*) bovine albumin fraction V, 0.2% (*w*/*v*) dextrose, and 0.3 % (*v*/*v*) catalase (beef).

### Susceptibility testing

Antibiotic susceptibility was assessed using the broth microdilution method for MIC determination. The effect of two combined inhibitors was determined using a checkerboard assay.(70) A liquid culture of *Msm* was incubated at 37 °C with aeration for 48 h, after which the culture had reached the stationary phase (Figure S11), and subsequently diluted in media containing diverse concentrations of the compound of interest (final concentrations of 0.25 - 128 µg/mL, total volume 200 µL) to OD_600_ 0.05 in a sterile 96-well plate (Corning Costar, flat-bottom, cell culture-treated). Positive controls consisted of samples containing only *Msm* culture in media and negative controls omitted *Msm* inoculation and contained only media. In line with reported procedures for *Msm* MIC assays,(33, 46, 71) we applied an incubation period of 4 days to all MIC experiments. Then, resazurin (20 µL of a 150 µg/mL stock solution) was added to all wells, and the samples were incubated at 37 °C for 12 h. The MIC was defined as the lowest concentration at which no resazurin colour change was observed. All assays were performed in triplicate.

### Fluorescent peptide incorporation assays

A 5 mL culture of *Msm* was grown at 37 °C to an OD_600_ of 0.8. This culture (196 µL) was added to a sterile 96-well plate (Corning Costar, flat-bottom, cell culture-treated). To this was added either TetraRh (2 µL, to a final concentration of 0.5 µM) or PentaFI (2 µL, to a final concentration of 5 µM), and varying concentrations of the compound of interest (2 µL, final concentrations ranging from 0.5 µg/mL to 256 µg/mL). A positive control (no inhibitor) and a negative control (100 µg/mL meropenem and 100 µg/mL clavulanic acid) were included. This was incubated at 37 °C for 24 h. Samples were then washed in PBS (200 µL, 3x), and fixed with 2% (*v/v*) in PBS for 30 min. The *Msm* cells were washed with PBS (200 µL, 3x), then resuspended in 200 µL PBS. The fluorescence of the *Msm* cells was analysed using an LSRFortessa™ X-20 (BD Biosciences). Cells treated with TetraRh were assessed using a 561 nm laser and a 586/15 bandpass filter. Cells treated with PentaFI were assessed using a 488 nm laser and a 530/30 bandpass filter. For each dataset 10,000 events were counted. Data were analysed using FlowJo™ Software. All assays were performed in duplicate.

### Biochemical inhibition assays

Ldt_Mt2_ dose-response assays were performed as described.(65) In brief, Ldt_Mt2_ (100 nM) was incubated with varying concentrations of the compound of interest (final concentrations ranging between 400 µM and 20.3 nM) for 10 min in the assay buffer (50 mM HEPES, pH 7.2, 0.01% (*v*/*v*) Triton X-100) and then assayed using Probe 1 (25 µM).

The ‘intrinsic’ thiol reactivity (*k*_chem_) was determined as described.(31) In brief, L-glutathione (500 nM) was incubated with varying concentrations of the compound of interest (final concentrations ranging between 400 µM and 20.3 nM) and Probe 1 (10 µM) for 15 h in assay buffer (50 mM HEPES, pH 7.2, 0.01% (*v*/*v*) Triton X-100).

The second-order rate constant of covalent target inactivation (*k*_inact_/K_I_) of Ldt_Mt2_ was determined as described.(31) In brief, Ldt_Mt2_ (100 nM) was incubated with varying concentrations of the compound of interest (final concentrations ranging between 400 µM and 20.3 nM) and Probe 1 (10 µM) for 3.5 h in assay buffer (50 mM HEPES, pH 7.2, 0.01% (*v*/*v*) Triton X-100).

Dose-response assays of PBP3 were assessed using the fluorometric S2d assay,(72–74) applying the procedure optimised for *P. aeruginosa* PBP3.(75) PBP3 (300 nM) in assay buffer (50 mM HEPES pH 7.4, 100 mM NaCl, 0.01% (*v*/*v*) Triton X-100) was incubated with varying concentrations of the compound of interest (final concentrations ranging between 400 µM and 20.3 nM) for 10 minutes in a black polystyrene, flat-bottomed 384-well μ-clear plate (clear bottomed, Greiner Bio-One, part number 781096). Then, a mixture containing S2d (1.5 mM), monobromobimane (mBBr; 0.05 mM), and D-Ala (1 mM) in assay buffer was added (final volume 25 µL). The fluorescence signal was measured using a BMG Labtech CLARIOstar instrument with λ_ex_=394 nm and λ_em_=490 nm. Data were analysed using Prism (GraphPad).

### Mass spectrometry assays

Protein-observed SPE-MS experiments with Ldt_Mt2_ were performed as described.(31) In brief, Ldt_Mt2_ (1 μM) in 50 mM tris, pH 7.5 was incubated with an inhibitor (100 µM) at room temperature. Mass spectrometry was performed using a RapidFire200 integrated autosampler/solid phase extraction (SPE) system (Agilent Technologies) employing a C4 cartridge (Agilent Technologies), coupled to an API40000 triple quadrupole mass spectrometer (Applied Biosystems) operating in the positive ionisation mode. The mass spectrometer parameters were: capillary voltage (2000 V), nozzle voltage (1500 V), fragmentor voltage (150 V), gas temperature (225 °C), gas flow (13 L/min), sheath gas temperature (300 °C), sheath gas flow (12 L/min).

## Author contributions

M.d.M., K.C.T., and C.J.S conceived the experiments; M.d.M. carried out experiments, with assistance from K.C.T., and P.R.; M.d.M. analysed the data; M.d.M. drafted the manuscript with C.J.S, with input from all authors.

## Acknowledgements

The project was co-funded by the Tres Cantos Open Lab Foundation (Project TC297). It was also supported by funding from the Biotechnology and Biological Sciences Research Council (BBSRC) [BB/M011224/1] and the Wellcome Trust (106244/Z/14/Z). P.R. thanks the Wellcome Trust (227298/Z/23/Z). We thank Robert Hedley and Vasiliki Tsioligka for providing technical assistance in the analysis of the fluorescent peptide incorporation assays at The Don Mason Facility of Flow Cytometry, Sir William Dunn School of Pathology, University of Oxford. We thank Alistair Farley, University of Oxford, for helpful discussions on the selection of sulfonyl pyridines.

